# Analyzing the implications of protein folding delay caused by translation

**DOI:** 10.1101/2024.01.27.577370

**Authors:** Bert Houben, Ramon Duran-Romaña, Paula Fernández Migens, Frederic Rousseau, Joost Schymkowitz

## Abstract

Because of vectorial protein production, residues that interact in the native protein structure but are distantly separated in the primary sequence are unavailable simultaneously. Instead, there is a temporal delay during which the N-terminal interaction partner is vulnerable to off-pathway, non-native interactions. In this analysis, we introduce “FoldDelay” (FD), a metric that integrates the topological pattern of atomic interactions of the native structure with translation kinetics to quantify such time delays. The FD metric reveals that many proteins, particularly at eukaryotic translation rates, exhibit residues with FDs in the range of tens of seconds. These residues, predominantly in well-structured, buried regions, often coincide with predicted aggregation-prone regions. We show a correlation between FD and co-translational engagement by the yeast Hsp70 chaperone Ssb, suggesting that fold-delayed regions have a propensity to misfold. In support of this, we show that proteins with high FDs are more frequently co-translationally ubiquitinated and prone to aggregate upon Ssb deletion. Finally, we find that FD cannot be adequately reduced through codon optimization, highlighting the importance of co-translational chaperones to shield these vulnerable regions. This work offers insights into co-translational proteostasis and the delicate balance between efficient folding and potential misfolding and aggregation during translation.

## INTRODUCTION

The functionality of globular proteins relies on adopting a three-dimensional shape known as the native structure. Achieving this structure involves folding an elongated polypeptide chain into a specific conformation. Although this process is intricate, it is widely acknowledged that all the necessary information for a protein to attain its native fold is encoded in its primary amino acid sequence [1]. Additionally, proteins can fold completely in physiologically relevant timescales, generally in the range of microseconds to seconds [2]. Most of these folding rates are derived from classic *in vitro* experiments in which the (re)folding of purified, full-length protein is monitored. These experiments yielded invaluable insights, including the realization that *in vitro* folding rates are partly determined by topological complexity [3].

However, protein translation is an aspect that is overlooked in such experiments. Protein translation progresses at an average rate of about 20 aas/s in prokaryotes and around five aas/s in eukaryotes [4–6], meaning that the complete synthesis of proteins can take seconds and even up to minutes. Hence, *in vivo*, local folding events often take place while a polypeptide chain emerges from the ribosome, i.e. co-translationally. Indeed, it is estimated that one-third of the *E. coli* cytosolic proteome folds at least one entire domain co-translationally [7], and this fraction is likely higher in eukaryotes, given their slower translation rates. The vectorial nature of translation is exploited *in vivo* to increase folding efficiency: the gradual addition of residues allows the growing polypeptide chain to sample stabilizing native interactions in a reduced conformation space, thereby avoiding kinetic traps associated with interactions with not yet formed residues towards the C-terminus in the sequence [8]. Indeed, some proteins fold more efficiently co-than post-translationally [9–13]. In line with this, co-translational folding has been put forth as one of the explanations for why a large portion of the *E. coli* cytoplasmic proteome does not reassemble back into their native folds after cell lysis and protein denaturation [14].

However, co-translational folding is a double-edged sword. While the incremental emergence of the nascent chain is beneficial, as it effectively promotes short-range native interactions in a reduced conformational space, the opposite is probably true for long-range interactors. Vectorial polypeptide production condemns residues with long-range interaction partners to idle while the remainder of the polypeptide chain is being produced, making them more vulnerable to non-native intra– and intermolecular interactions that can potentially lead to “premature folding” – i.e. misfolding – and/or aggregation. In support of this, several sources report that newly synthesized proteins are more vulnerable to misfolding and aggregation than existing, matured proteins [15, 16], with topologically complex proteins being more at risk [16]. It is also widely understood that premature co-translational misfolding – before a critical nascent chain length is achieved – is indeed detrimental and must be avoided [17]. Proteins have, therefore, evolved to prevent premature intermolecular interactions. Natan et al., for example, show a systematic enrichment of multimerization domains near the C-terminus of proteins, whereas the artificial placement of those domains at the N-terminus gives rise to misfolding and aggregation [18]. To avoid premature co-translational misfolding events, an entire branch of the proteostasis network (PN) exists that acts specifically at the translation stage, shielding vulnerable regions. Firstly, ribosomes themselves have a holdase function as their negatively charged surface interacts with nascent chains, preferentially with basic and aromatic residues, thereby preventing premature co-translational folding by destabilizing the nascent chain [17, 19, 20]. Secondly, a host of dedicated chaperones engage nascent chains at the ribosome [21]. The typical example of this is Trigger Factor (TF) in *E. coli,* which directly interacts with both the nascent chain and the ribosome, thereby preventing off-pathway interactions [22]. In eukaryotes, co-translational chaperones are most well-studied in *S. cerevisiae,* in which Nascent polypeptide Associated Complex (NAC) and Ribosome Associated Complex (RAC) directly engage the ribosome and interact with the nascent chain near the ribosome exit tunnel [21]. RAC recruits an Hsp70-type ribosome-associated chaperone called Ssb, which prevents premature folding through binding-release cycles [23–25]. Still, this mechanism is not foolproof as an estimated one-third of newly synthesized polypeptides are targeted for proteasomal degradation, either through mistakes in translation or inability to attain the native fold [26].

Clearly, the vectorial nature of translation can benefit folding outcomes, but it also poses a risk as residues that idle for extended periods of time on the ribosome can potentially engage in off-pathway interactions, necessitating a dedicated co-translational proteostasis network. In this work, we describe a method to identify vulnerable regions in co-translational folding. To do this, we quantify the length of time between the translation of each residue and that of all its native interaction partners. Effectively, we calculate the delay on co-translational folding experienced by individual residues, for which we coin the term “FoldDelay” (FD). Using the FD algorithm, we show that many proteins contain residues with FDs in the range of minutes, especially at eukaryotic translation rates. Furthermore, we establish that residues with the strongest FDs tend to be in well-structured, buried parts of globular proteins and are often part of predicted aggregation-prone regions (APRs). In addition, we show that *in vivo*, the yeast co-translational Hsp70 chaperone Ssb preferentially engages fold-delayed regions. Aggregation propensity in these fold-delayed Ssb binding sites causes co-translational aggregation upon Ssb knockout. Both these findings suggest that regions of high FD are indeed at high risk of co-translational misfolding and aggregation. In support of this, we further show that proteins with high FDs are more frequently co-translationally ubiquitinated. Finally, we address whether FD was mitigated evolutionarily through codon optimization. We conclude that this is an unlikely evolutionary strategy, and it is probably the co-evolution with co-translational chaperones that reduced the risk of increased FD associated with topologically complex proteins.

## RESULTS

### Protein translation is orders of magnitude slower than protein folding

To visualize the differences in timescales of protein folding and translation rates, we directly compare folding rates and estimated translation times for 133 proteins that have experimentally recorded *in vitro* refolding rates from denaturing conditions reported in the Protein Folding DataBase (PFDB) [27] (**Figure 1A**). The PFDB contains information on proteins from a wide array of species, and for a lot of these, an accurate translation rate has never been established. Therefore, translation times were estimated by multiplying protein lengths with an average translation rate. We assumed a relatively fast translation rate of 20 aas/s for all prokaryotic proteins and five aas/s for all eukaryotic proteins [4–6].

**Figure 1.**
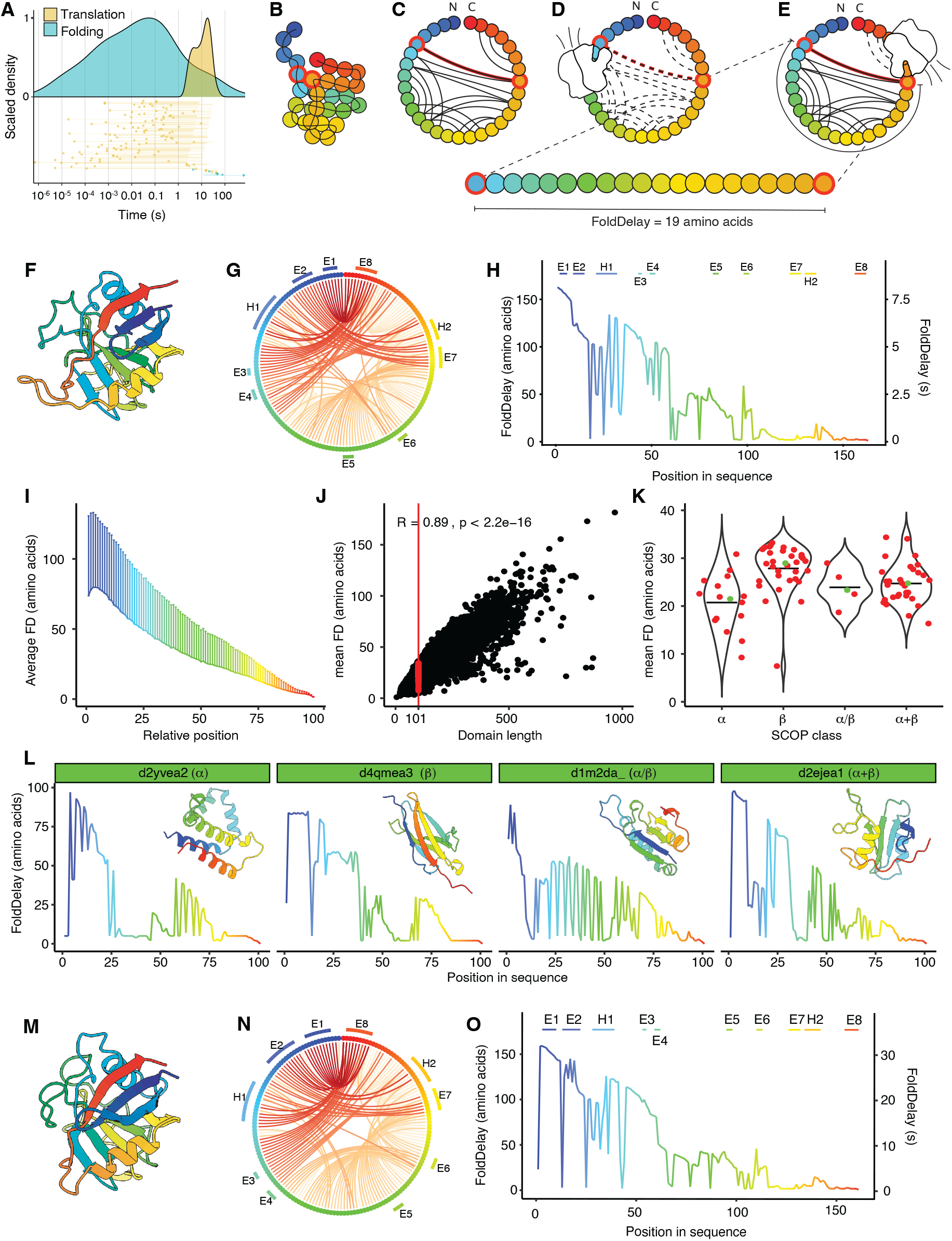
– FoldDelay measures waiting times incurred by nascent residues during translation. (**A**) Distribution of average folding times versus estimated average translation times of 133 proteins in the Protein Folding Database (PFD2.0 [27]). Arrows show difference in folding and translation times for individual proteins. Yellow arrows indicate proteins for which average translation time is slower than average folding time, blue arrows indicate proteins for which average translation time is faster than average folding time. **(B)** Schematic representation of the globular native structure of a hypothetical protein. Amino acids are colored in a gradient from N-term (blue) to C-term (red). **(C)** Contact map of hypothetical structure in (A), with contacts indicated by solid lines. **(D)** Contact map as residue 5 emerges from the ribosome. Dotted lines indicate interactions that are not yet accessible as not all interaction partners have been added to the polypeptide chain. **(E)** Contact map as residue 24 emerges from the ribosome. Solid lines indicate interactions that are now available, dotted lines indicate interactions that are not. All contacts for residue 5 have at this point become available. The FD incurred by residue 5 is 19 amino acids, spanning the point where residue 5 emerged from the ribosome, until the point where its most C-terminal interaction partner, residue 24, emerges. **(F)** Cartoon representation of the native structure of the *E. coli* peptidyl-prolyl isomerase B (PPI B, UniProt code P23869) enzyme as predicted by AlphaFold. Residues are colored on a gradient from N-term (blue) to C-term (red). **(G)** Contact map of PPI B from the structure in (F). **(H)** Per-residue FD calculation for PPI B. **(I)** Mean FD of domains in the SCOPe40 dataset versus the relative residue position in the domain (scaled from 1 to 100). Error bars indicate standard deviation. **(J)** Domain length versus mean FD of the SCOPe40 dataset. Red points indicate domains of exactly 101 amino acids, the domain length with the most datapoints in the SCOPe40 database. **(K)** Violin plots showing the distribution of mean FD for the domains of exactly 101 amino acids per SCOP class. Green points indicate a representative example in each group (domains with mean FD closest to the median of their respective SCOP class). **(L)** FD profiles of the representative examples for each SCOP class indicated in (K). (**M)** Cartoon representation of the native structure of a peptidyl prolyl cis-trans isomerase from *S. cerevisiae* (UniProt code P14832) as predicted by AlphaFold. Residues are colored on a gradient from N-term (blue) to C-term (red). **(N)** Contact map of the structure in (F). **(O)** Per-residue FD calculation for the structure in (F).

Despite this, the distributions of translation times and folding times are clearly separated (**Figure 1A**). Translation times are typically on the order of seconds, whereas folding times range from microseconds to seconds, and in 126 of 133 cases (95%), the *in vitro* refolding time of the full-length protein is shorter than the time estimated to complete its translation (**Figure 1A**). Moreover, we here consider the time it takes for an entire polypeptide chain to cooperatively fold to its native conformation. Local protein conformational dynamics are generally even faster, ranging from nanoseconds to microseconds [28]. As a result, for many proteins folding is a co-translational process that starts as soon as the N-terminal part of the protein emerges from the ribosome tunnel and long before the full-length protein chain has been synthesized and released from the ribosome.

### Vectorial protein translation imposes spatial and temporal constraints on folding

Protein folding studies, both *in vitro* and in the cell, have demonstrated that protein topology (i.e. the sequence order in which the structural elements of the tertiary fold occur in the primary sequence) is a key determinant of folding rates and efficiencies [29]. Protein topological complexity is often described by Contact Order (CO), a metric which calculates the average sequence distance separation of native interactions.

As shown in **Figure 1A**, translation is relatively slow compared to folding. This means that during protein translation topology restricts folding not only spatially but also temporally: residues that are separated spatially in the primary sequence are also separated temporally as interaction partners towards the C-terminal end of the protein will simply not exist until they have been translated. Inspired by and building on CO, we here propose a metric that accounts for these temporal topological constraints, for which we have coined the term “FoldDelay” (FD). For each residue (*i*) in a protein sequence, FD measures the sequence distance (ΔS) from its furthest away C-terminal interactor (*j*):

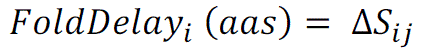

In other words, FD measures the number of residues that need to be synthesized before residue (*i*) can engage with all its native interaction partners. The assumption is that until that moment, its folding is delayed (hence “FoldDelay”). FD can also be expressed in time units by factoring the decoding times (*t_dec_*) of the different residues between *i* and *j*:

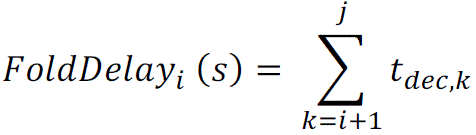

The FD calculation is schematically represented in **Figures 1B-E**. **Figure 1B** shows the native fold for a hypothetical small globular protein. From this native structure, all interactions are mapped, resulting in a contact map (**Figure 1C**). While in post-translational folding all these contacts are available simultaneously (**Figure 1C**), in the co-translational paradigm the contact map changes over time (**Figure 1D and E)**. Residue 5, for example, interacts with residue 24 in the native structure (residues outlined in red in panels **Figures 1B-E)**. As residue 5 emerges from the ribosome, this interaction is not available, as residue 24 has not been added to the polypeptide chain (**Figure 1D**). Therefore, residue 5 cannot form all its native interactions until residue 24 has emerged from the ribosome and becomes physically accessible (**Figure 1E**). Residue 5, therefore, incurs a FD of 19 aas. Importantly, residue 24 has practically no FD as all its long-range interaction partners (residues 5 and 6) have already been added to the polypeptide when it emerges from the ribosome exit tunnel.

As an example, **Figures 1F-H** show the FD calculation for the *E. coli* peptidyl-prolyl isomerase B (PPI B, UniProt code P23869) protein. PPI B is an abundant cytoplasmic enzyme with a length of 164 amino acids. Its functional form is a globular shape comprised of beta sheets and alpha helices separated by several random coils (**Figure 1F**). While PPI B has an average folding time of about 600 μsits estimated translation time is eight seconds (assuming an average translation rate of 20 aas/s). Clearly, the timescales of folding and translation here are vastly different, and PPI B is likely to start folding co-translationally. **Figures 1G and 1H** show the contact map calculated from the PPI B native structure and the per-residue FD profile, respectively. PPI B contains a beta-sheet comprised of a strand close to the N-terminus (E1), and a strand close to the C-terminal end in the primary sequence (E8). Strand E1 cannot fully be stabilized in its native conformation until strand E8 has been produced, which takes about 160 aas. This means that strand E1 sits partially exposed for about eight full seconds. On the other hand, strand E8 has a negligible FD, as its interaction partners have all been produced when it emerges from the ribosome.

To explore general FD patterns across different protein topologies, we ran FD analyses on the protein domains of the SCOPe40 dataset. This dataset contains single-domain structures that have been manually classified based on their architectures and filtered so that no two domains in the set have more than 40% identical sequences [30, 31]. Reflecting the vectorial nature of protein translation, FD has both spatial and temporal implications. First, the FD profiles of proteins display an N-to C-terminal gradient: N-terminal elements generally incur larger FDs than more C-terminal elements **(Figure 1I)**. In addition, domain size is a big determinant of FD, as the longer a polypeptide chain, the more potential there is for long-range interactions, leading to high FDs (**Figure 1J**). On top of length, FD also reflects the topological specificity of the translated protein. Indeed, even when considering proteins of identical length (101 aas), proteins from different SCOPe classifications have different FD patterns (**Figure 1J and K**). More complex protein topologies have higher FDs and present more pronounced N– to C-terminal FD gradients, resulting in different profiles for alpha-helical or beta-sheet structured domains (**Figure 1K and L**). Interestingly, FD profiles of large multi-domain proteins often display a sawtooth profile reflecting the domain dependence of N– to C-terminal FD gradients (**figure S1**).

On top of topology, FD is also dependent on translation rates, which can vary strongly between species. **Figure 1M** shows the AlphaFold predicted structure of a peptidyl-prolyl cis-trans isomerase (CPR1, UniProt code P14832) from *S. cerevisiae*, which is homologous and structurally very similar to PPI B from *E. coli.* As is the case for PPI B, the N-terminal domain has a strand near its N-terminus (E1) that forms contacts with a strand at the C-terminus of the domain (E8), resulting in high FD values for E1 (**Figure 1N and O**). Although the FDs for the N-terminal strands in PPI B and CPR1 are similar when expressed in number of aas, the relatively slower translation rates of *S. cerevisiae* (estimated to be around 5 aas/s on average), means that strand E1 idles on the ribosome for about 32 seconds, as opposed to the 8 seconds estimated for strand E1 in PPI B. Therefore, differences in translation rates of different organisms can cause domains with very similar folds to incur vastly different FDs.

### Sequence segments with high FoldDelays often consist of aggregation-prone tertiary structural elements that stabilize the native structure

Having established the FD algorithm, we next used it to explore FD patterns on a proteome-wide scale. The near-exhaustive availability of AlphaFold-predicted structures combined with the computationally inexpensive nature of our algorithm allows us to calculate FD for all residues across entire proteomes [32, 33]. In addition, AlphaFold models provide a confidence measurement to assess the relative position of two residues within the predicted structure, called the Predicted Aligned Error (PAE). We used this metric to filter out interactions between residues whose relative positions with respect to each other are predicted with low confidence since these interactions most probably do not occur in the actual structure, as is the case for contacts with disordered regions or some contacts between distinct domains. (**figure S2**).

We calculated the FD incurred by all residues in the *E. coli* and *S. cerevisiae* proteomes, assuming average translation rates of 20 aa/s and 5 aas/s, respectively. Interestingly, most proteins have at least one residue that has to wait for tens of seconds for the translation of all its native interacting residues (**Figure 2A**). Binning proteome-wide FD however, reveals that most residues have low FDs as they interact only with their neighbors (± 5 aa). While intermediate FDs are relatively rare, about 23% of residues in *S. cerevisiae* proteome incur FDs of more than 10 seconds (**Figure 2B**), while the same is true for 7% of *E. coli* residues (**figure S3A**). Specific secondary structures are more likely to incur FD (**Figure 2C and figure S3B**). Logically, residues in random coils (C) are depleted from the high FD groups as they barely make any contacts. On the other hand, helical structures (G, H and I) are dominated by short-range contacts, yielding average FDs. Pi-helices (I) have higher FDs than alpha-helices (H), which makes sense given that backbone interactions in pi-helices occur at an interval of five residues, where this is four residues for alpha-helices and three for 3-turn helices (G). Finally, beta-structured elements (B, E), are enriched in residues with the highest FDs. Again, this makes sense given that contacts between beta strands are generally more long-range than those between residues in alpha helices.

**Figure 2.**
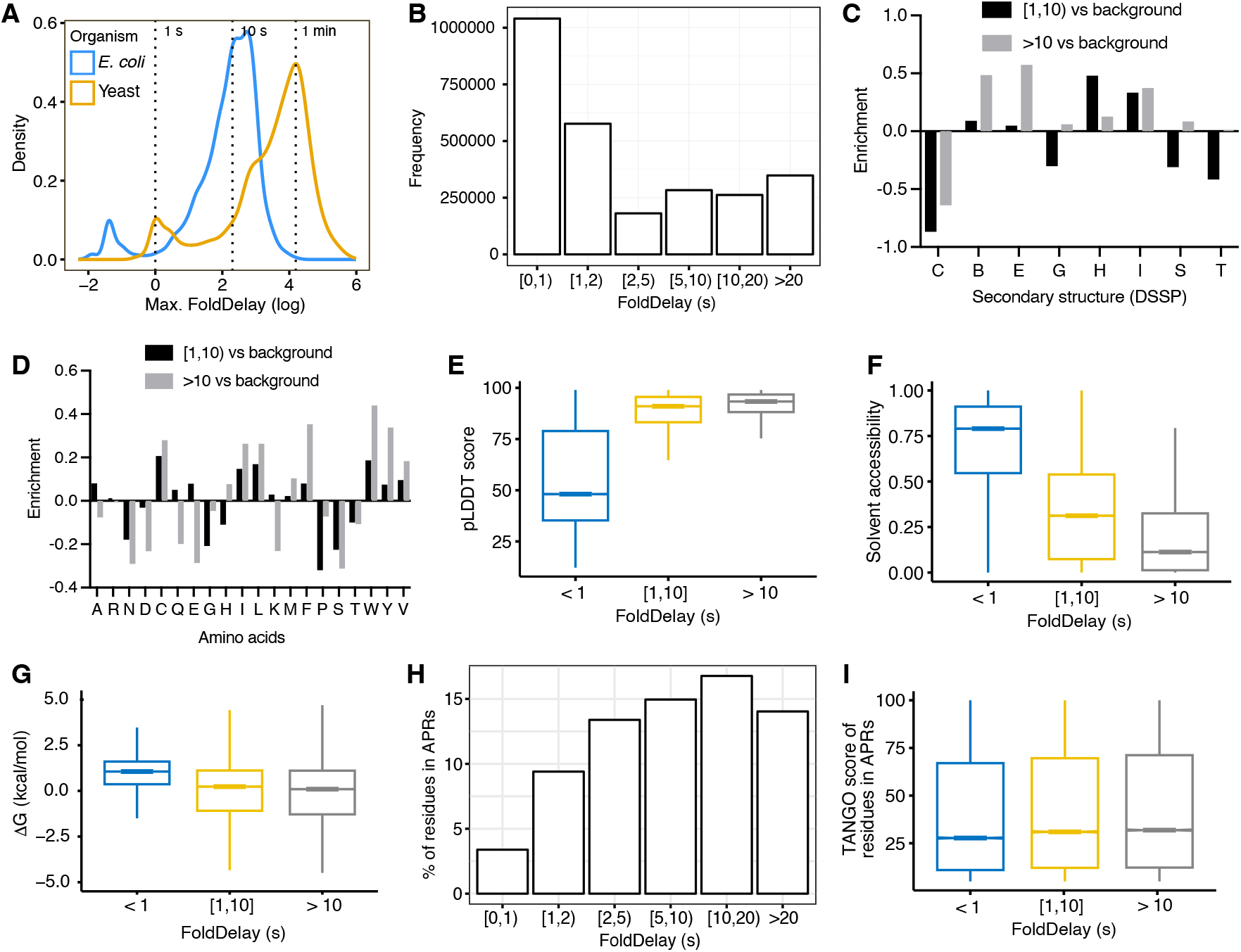
– Exploring FoldDelay across proteomes. (**A**) Distribution of residues with the maximum FD (log scale) for each protein in *E. coli* (n = 3,910) and *S. cerevisiae* (yeast; n = 5,812). Vertical dotted lines indicate 1 second, 10 seconds and 1 minute. **(B)** Number of residues for different FD bins in yeast proteins. **(C)** Enrichment of residues with FDs between 1 and 10 seconds (or bigger than 10 seconds) versus background for the different DSSP secondary structure categories in yeast proteins. C = coil, B = β-bridge, E = extended strand in β-sheet conformation, G= 3-turn helix, H = 4-turn helix, I = 5-turn helix, S = bend and T = hydrogen bounded turn. **(D)** Enrichment of residues with FD between 1 and 10 seconds (or bigger than 10 seconds) versus background for all amino acid types in yeast proteins. **(E-G)** pLDDT scores (E), solvent accessibilities (F) and stabilities for residues in yeast proteins for different categories of FD. **(H)** Percentage of residues in APRs (TANGO score > 5) for different FD bins in yeast proteins. To avoid biases, residues in transmembrane domains and signal peptides were filtered out. **(I)** Aggregation strength (TANGO score) for residues in APRs of yeast proteins for different categories of FD.

Looking at the sequence composition of segments with high FDs, we find them to be enriched in aromatic and aliphatic residues (**Figure 2D and figure S3C**). This makes sense as these residues are often buried in the hydrophobic cores of globular proteins, where they make many contacts. Exploring this further, we find that regions of high FD are often structurally ordered – as indicated by the AlphaFold pLDDT score, which inversely correlates with disorder – (**Figure 2E and figure S3D**) and indeed constituted of buried residues – as shown by their relatively low solvent accessibility (**Figure 2F and figure S3E**). Furthermore, regions of high FD are usually stabilizing to the domain structure, as shown by their low free energy (**Figure 2G and figure S3F**). Given their propensity for beta-sheet formation and hydrophobic nature, we asked whether regions of high FD tend to be aggregation-prone. Indeed, we find that the proportion of residues in aggregation-prone regions (APRs) substantially increases with FD (**Figure 2H and figure S3G**), although the distribution of their aggregation propensities is quite similar (**Figure 2I and figure S3H**).

### The co-translational chaperone Ssb binds to regions with high FoldDelays

Aggregation-prone exposed regions of high hydrophobicity are the preferred binding sites of many molecular chaperones, including Hsp70s [24, 34, 35]. It is proposed that Hsp70s bind to these regions to delay the folding of newly forming polypeptides until the residues required for folding emerge from the ribosome, thus preventing the formation of non-native interactions [35, 36]. Given that FD reflects co-translational exposure and that regions of high FD tend to be hydrophobic, we hypothesized that FD could help explain the engagement of specific segments of the nascent chain by chaperones. To address this question, we used a dataset containing the binding footprints for the co-translational chaperone Ssb from *S. cerevisiae,* obtained by Döring *et al*. [24] using selective ribosome profiling (SeRP). These binding footprints indicate the codons that are being translated by ribosomes while Ssb is bound to the emerging polypeptide chain (**Figure 3A**).

**Figure 3.**
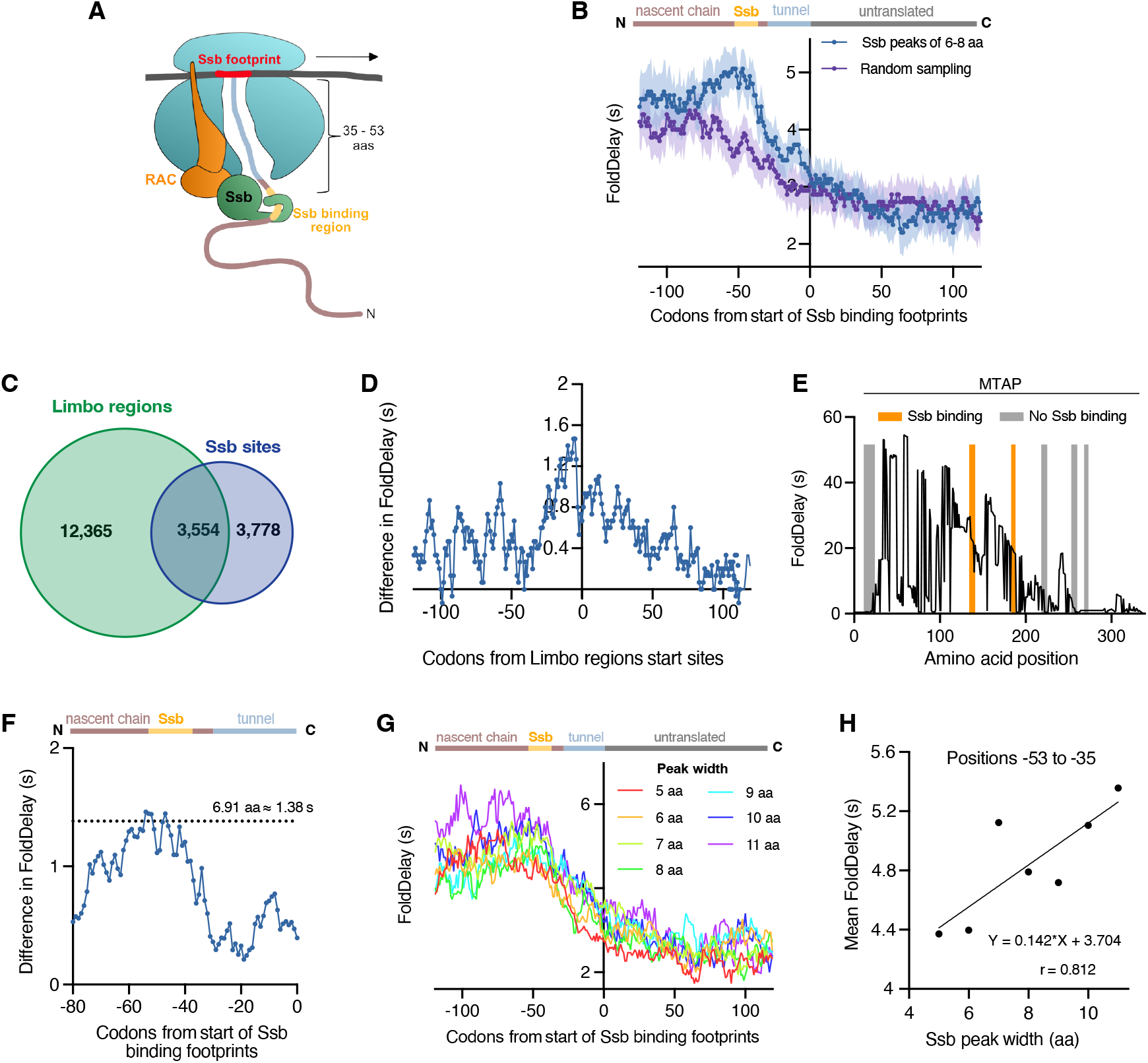
– Ssb binds to regions with high FoldDelays. (**A**) Schematic representation of Ssb with a nascent chain during translation. Ssb is targeted to the nascent chain by the ribosome-associated complex (RAC) once the nascent chain reaches a length of around 50 aas. **(B)** FD in the nascent chain at the start of Ssb binding for sites with a peak width between 6-8 aas (n = 3,371), as compared to an equivalent number of randomly sampled positions (n = 4,000) from the same set of proteins. The line represents the median value at each position, while the shaded region is the 95% bootstrapped confidence interval (CI). **(C)** Overlap between limbo regions and Ssb binding sites (width of 5-11 aas) (**D**) Difference between the median FD values of Limbo regions that are also Ssb binding sites, and Limbo regions that are not Ssb binding sites. **(E)** FD profile of MTAP. Limbo regions that are also Ssb binding sites are shown in orange while those that are not Ssb binding sites are shown in grey. **(F)** Difference between the median FD values of the aligned Ssb footprints and of the randomly sampled positions showed in B. Dotted line indicates the average peak width of Ssb binding sites in the dataset. **(G)** FD in the nascent chain at the start of Ssb binding for sites with a peak width of 5 aa (n = 1,412), 6 aa (n= 1,277), 7 aa (n = 1,111), 8 aa (n = 983), 9 aa (n = 945), 10 aa (n = 897) and 11 aa (n = 707). **(H)** Average median FDs between positions –53 and –35 (Ssb binding region) per width peak of aligned Ssb binding footprints with a peak width between 5-11 aas. Based on the linear model the average FD value at these positions increases with Ssb peak width. All experimental Ssb binding sites used in this figure are derived from [24].

We carried out a metagene analysis by aligning the starting site of Ssb binding footprints across the *S. cerevisiae* proteome and calculated the median FD value at each position. A distinct FD peak was revealed at around 50 aa towards the N-terminal side (**Figure 3B**). This is the exact distance that has been reported to exist between the Ssb footprint, i.e., the sequence segment protected by the ribosome at the moment of Ssb engaging the nascent chain, and the actual Ssb binding site [24, 37]. This indicates that the observed FD peak is directly associated with the regions engaged by Ssb. Indeed, at these positions, we observed some of the characteristic sequence and structural properties of Ssb binding motifs [24, 25], including an enrichment in positively charged residues and β-sheet propensity (**figures S4A and S4B**), and an underrepresentation of intrinsically disordered regions (**figure S4C**). A similar FD pattern was observed using a different published dataset of Ssb binding regions [25] (**figure S4D**). On the other hand, a dataset of Ssb binding regions generated in the absence of RAC (*RAC*Δ, [24]), a cochaperone that is required for high affinity binding of Ssb to its substrates, did not show any peak around these positions (**figure S4E**). Interestingly, there is an additional FD peak between –16 and –6 aa from the start of Ssb biding footprints, which is approximately 36 residues downstream of the main Ssb binding region (**Figure 3B**). This peak might correspond to other Ssb binding regions, as these have been previously described to occur in proteins every 36 amino acids, on average [38]. Together, these results indicate that Ssb binds to regions with high FD.

Intriguingly, despite Ssb recognition motifs being extremely common within protein sequences [38], SeRP data showed that many putative binding sites *in vitro* are actually ignored *in vivo* [24, 25]. Thus, we investigated whether putative chaperone binding motifs with low FDs could be skipped co-transitionally by Ssb. To investigate this, we used the computational tool Limbo to predict chaperone binding sites in yeast proteins [39]. Although Limbo was trained to predict *E. coli* DnaK binding sites, these motifs have been shown to be very similar to Ssb binding regions [24]. In fact, Limbo regions are enriched around 50 residues upstream of Döring *et al*. [24] Ssb footprints (**figure S3F**) and match with half of the identified Ssb binding regions (**Figure 3C**). On the other hand, only 22% of all Limbo predicted regions matched with an Ssb binding region (*P*-value < 0.001 by Fisher exact test), suggesting that there is a higher level of regulation beyond the amino acid sequence. Comparing Limbo regions that matched and did not match with Ssb binding regions, we saw that those that are not bound by Ssb co-translationally have lower FDs, even when analyzing regions with similar relative positions in both groups to avoid biases (**Figure 3D and figures S4G and S4H**). As an example, the protein S-methyl-5’-thioadenosine phosphorylase (MTAP) has six predicted chaperone binding sites based on Limbo (**Figures 3E**). Out of these, only two were experimentally identified *in vivo* and reside in regions with high FDs. Conversely, the other four predicted binding sites are in regions with lower or even negligible FDs. It seems then that Ssb not only engages target based on amino acid composition, but also on availability, which is aptly captured by the FD metric. The experimentally determined Ssb binding sites in MTAP have a FD of about 20 seconds during which time they are available for Ssb engagement. The more C-terminal Ssb binding sites, on the other hand, have no FD as all their interacting residues have already been translated.

As discussed by Döring et al. in the original Ssb SeRP publication, the maximal lifetime of the Ssb-Nascent chain complex can be extrapolated from the width of Ssb-binding peaks [24]. The average width of the Ssb peaks we consider in our analysis is 6.9 aas, which corresponds to a translation time, and hence Ssb engagement time, of 1.38 seconds. Intriguingly, we found that FDs of experimentally confirmed Ssb binding regions are, on average, 1.44 s higher FDs than regions from the same proteins that were sampled at random (**Figure 3G**). This suggests that regions that have a FD that is equal to or higher than the Ssb binding time can actually be engaged by the chaperone. To corroborate this, we asked whether Ssb binding sites with longer engagement times, i.e., wider footprints, have higher FDs. For Ssb footprints ranging in size from 5 to 11 aas, we indeed observed a strong positive correlation between the FD values at positions –53 to –35 (Ssb binding region) and the footprint size (*p*-value = 0.04) (**Figures 3G and 3H**). Moreover, the slope of this correlation roughly corresponds to the addition of one amino acid (**Figure 3H**). Correlations outside the Ssb binding region were weaker and not significant (**figures S4I and S4J**).

### Proteins with high FoldDelays are associated with co-translational misfolding and aggregation

We have shown that Ssb preferentially engages regions of high FD. To corroborate this, we used a dataset produced by Willmund et al., who mapped Ssb clients across the *S. cerevisiae* proteome and showed that the deletion of Ssb leads to widespread aggregation of newly synthesized polypeptides [23]. We used this dataset to assess whether Ssb clients indeed have higher FDs and whether proteins with high FDs are disproportionately affected by Ssb deletion. To this end, we assigned a single value to each protein by simply summing the FDs of individual residues. As expected, Ssb clients generally have higher total FDs than proteins that are not engaged by the co-translational chaperone (**Figure 4A**). Furthermore, Ssb clients that aggregate upon deletion of Ssb (*SSB*Δ, [23]) have a higher total FD than Ssb substrates that remain soluble (**Figure 4A**). To examine this difference in more detail, we looked at the metagene FD profile of specific Ssb binding sites of aggregated and soluble Ssb substrates based on Döring *et al*. [24] ribosome footprints. Ssb binding regions in proteins that aggregate in *SSB*Δ cells have, on average, a one second higher FD compared to binding regions in proteins that do not aggregate (**Figure 4B and figures S5A and S5B).** We next investigated whether proteins in the aggregated fraction upon Ssb deletion have higher intrinsic aggregation propensities. Although these proteins have a similar number of APRs per length unit (**Figure 4C),** we found that proteins that aggregate upon Ssb deletion have a significantly higher proportion of APRs in their Ssb binding regions (positions –53 to –35) compared to other regions in the same proteins of the same size (**Figure 4D)**. In contrast, proteins that remain soluble have a significantly lower proportion of APRs in their Ssb binding regions (*P*-value < 0.0001 by Fisher exact test), similarly to other regions from the same proteins (**Figure 4E)**. This suggests that APRs are driving the aggregation of the aggregated proteins in *SSB*Δ cells.

**Figure 4.**
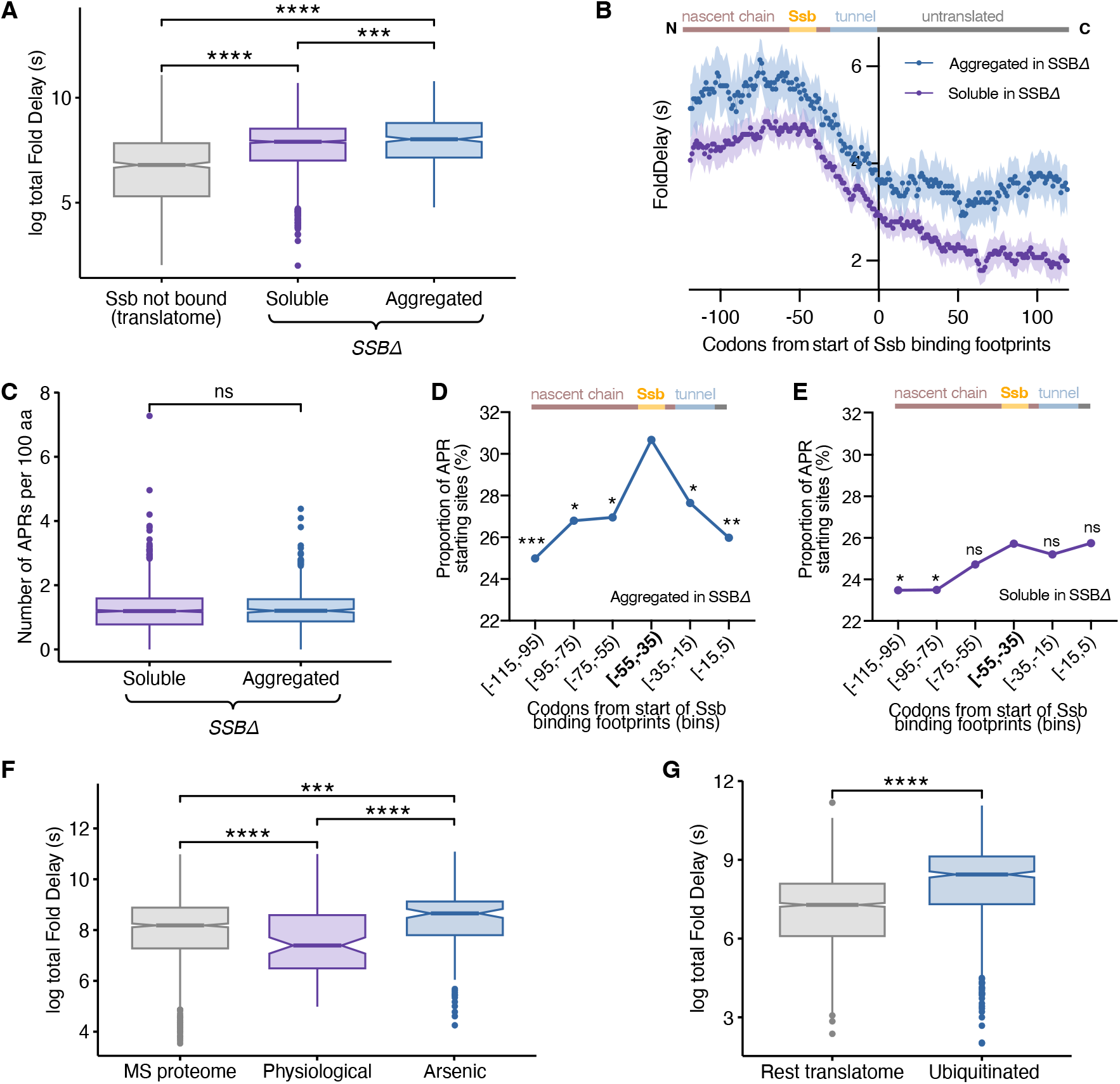
– Proteins with high FoldDelays are associated with co-translational misfolding and aggregation. (**A**) Total FD of proteins that are actively translated in *S. cerevisiae* (translatome), which have been stratified based on whether they interact co-translationally with Ssb (n = 1,913) or not (n = 910) [23]. Proteins that interact with Ssb are further stratified on whether they remain soluble (n = 1,495) or aggregate (n = 418) in *SSB*Δ cells. **(B)** FD in the nascent chain at the start of Ssb binding for sites with a peak width between 5-11 aas in proteins that aggregate or remain soluble in *SSB*Δ cells. There are 1,917 and 5,415 Ssb binding sites in proteins that aggregate or remain soluble, respectively. The line represents the median value at each position, while the shaded region is the 95% bootstrapped CI. (**C)** Number of APRs per 100aa in proteins bound by Ssb that remain soluble (n = 1,495) or aggregate (n = 418) in *SSB*Δ cells. **(D,E)** Percentage of APR starting sites in bins of equal length based on the start of Ssb binding footprints with a width between 5-11 aas in proteins that aggregate (D) or remain soluble (E) in *SSB*Δ cells. Fisher exact test with FDR correction was used to compare the proportion of APR starting sites at bin –55 to –35 (Ssb binding region) against the other bins. **(F)** Total FD of yeast proteins under physiological conditions (n = 107) or upon exposure to arsenite stress (n = 140) compared to background (MS proteome; n = 1,179) [40]. **(G)** Total FD of proteins that are co-translationally ubiquitinated under physiological conditions (n = 600) [43]. As background we use the translatome (n = 1,790) reported by Willmund et at [23]. Statistical significance was determined by unpaired Wilcoxon test with Bonferroni correction for multiple comparisons (A, C, F and G).

To further corroborate these findings, we analyzed a dataset produced by Jacobson et al. who identified proteins that aggregate upon treatment of yeast cells with arsenite [40], a metalloid known to cause aggregation by interfering with the folding of nascent proteins [41]. Again, we found that arsenite stress disproportionately causes the aggregation of proteins with high total FD (**Figure 4F**). In eukaryotic cells, misfolded proteins are tagged through ubiquitination for degradation [42]. Duttler *et al.* [43] showed that a subset of cytoplasmic nascent polypeptides is often co-translationally ubiquitinated. Re-analysis of this dataset, revealed that proteins that are co-translationally ubiquitinated have a significantly higher total FD compared to other abundantly translated yeast proteins (**Figure 4G**).

### FoldDelay cannot be fully compensated for through codon optimization

Our previous analyses suggest that fold-delayed regions are weak spots in co-translational folding that can jeopardize folding outcomes. We therefore asked whether genetic sequences are in any way optimized to reduce FD. In our analyses so far, we made use of flat translation rates across transcripts to estimate FDs in actual time units. However, during translation *in vivo,* codons are not translated at a flat rate. Instead, ribosome profiling studies revealed a variety in codon translation rates, even between codons encoding the same amino acid [44]. This means that FD can be reduced by preferentially using fast-translating, “optimal” codons in regions that span long-range interactions. To test this, we attempted to find correlations between FD and several codon optimization metrics, including the Codon Adaptation Index and the more recently described %MinMax [45]. However, we were unable to find convincing evidence of codon optimization towards reducing FD (data not shown).

We next asked what the actual reduction in FD would be, given perfect codon optimization. Using the typical codon translation rate tables for *S. cerevisiae* determined by Dana and Tuller [44] and Sharma et al [46] (**Figure 5A and E**, respectively), we recalculated the FD of the longest-range interaction in each *S. cerevisiae* protein using either the wild-type codon, yielding the “actual” FD or the synonymous codon with the fastest typical translation rate, giving the “minimal” FD (**Figure 5B and F**). In effect, the minimal translation rates represent the idealized situation where every codon between long-range interactors is optimized for speed. We then calculated the hypothetical time gained by fully optimizing sequences between long-range interactors (**Figure 5C and G**). Even at very high FDs, these gains seem to be marginal. Indeed, calculating the proportional reduction of FD given perfect optimization, we find that the actual FD could only be reduced by about 20% according to the Dana & Tuller decoding timetables, and around 15% according to the tables devised by Sharma et al. (**Figure 5D and H).** The reason we were unable to find codon optimization towards FD may, therefore, be that there is not much to gain. Moreover, optimizing long stretches of amino acids between interactors is evolutionarily a tall order, given that individual point mutations will have extremely marginal FD effects and are, therefore, unlikely to persist.

**Figure 5.**
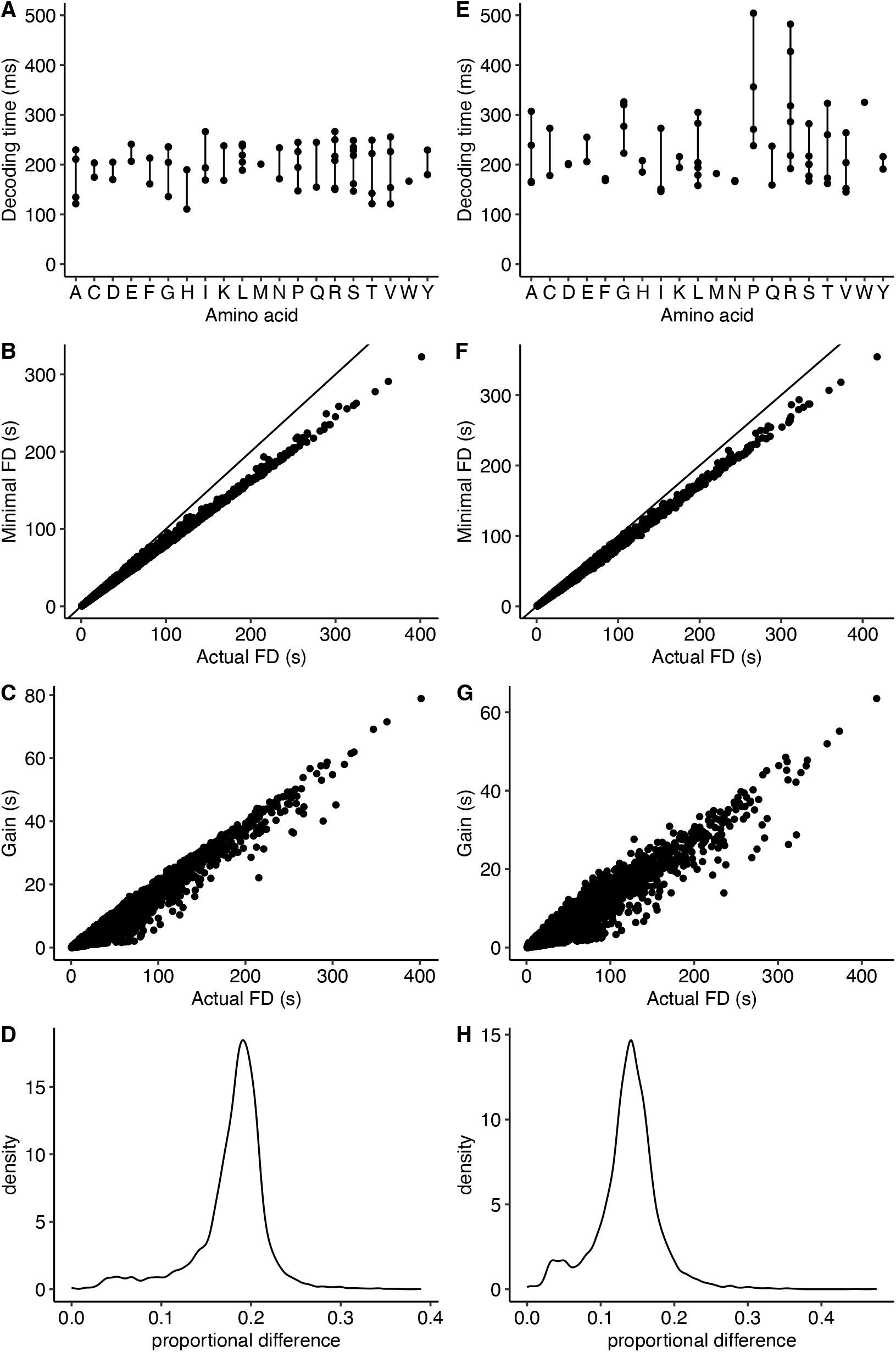
– Compensating FoldDelay through codon optimization is an implausible evolutionary strategy. (**A**) Distribution of codon decoding times per amino acid as reported by Tuller et al [44]. **(B)** Minimal FD vs. actual FD calculated per protein. For each protein’s furthest distance interaction, the actual and minimal translation times of the separating residues were calculated using the mean translation rates of the actual codons, and the mean translation rate for the fastest synonymous codon, respectively. **(C)** Gain in FD as calculated by the difference between the actual FD and the minimal FD in **(B)**. **(D)** Distribution of the proportional differences, calculated as the ratio between the differences shown in **(C)** and the actual FD. **(E-H)** Identical analyses as those shown in **(A-D)**, this time using the average decoding times reported by [46] based on data from [61].

## DISCUSSION

The work presented here describes FD, a metric that quantifies the time nascent residues idle on the ribosome before their native interaction partners are physically available. Conceptually, FD resembles the simple yet widely used and validated CO metric in that it captures topological complexity by the separation between interacting residues in the native structure [3]. However, in contrast to CO, FD measures the temporal separation caused by the translation process. Hence, FD can be considered an extension of CO that is more informative for the co-translational paradigm.

In recent years, it has become clear that co-translational folding is a common folding mechanism across proteomes and is often beneficial to folding outcomes [17]. FD, however, lays bare a potential shortcoming of this folding mechanism: the nascent chain becoming available incrementally creates a delay between translation and folding. Although this was hitherto implicitly understood in the co-translational folding field, we provide a systematic quantification of this phenomenon and show that this delay can, in fact, last for tens of seconds and even minutes (**Figure 2A and B**), a vast period on a molecular timescale [47]. During this time, such segments are potentially vulnerable to non-native interactions such as misfolding and aggregation. Moreover, we have shown that the regions with the highest FDs often occur in structured, hydrophobic regions that are meant to be buried within the hydrophobic core of the structure of globular proteins (**Figure 2 C-F**). Exactly such regions are likely to engage in homotypic off-pathway interactions leading to aggregation (**Figure 2H**). This begs the question of whether fold-delayed APRs could aggregate co-translationally, especially in the context of polysomes where there is a higher local concentration of nascent polypeptides exposing identical APRs. Supporting the hypothesis that aggregation may already occur during translation, a recent study found that mRNAs coding for protein constituents of insoluble aggregates are inherent components of those aggregates in the brain tissue of Alzheimer’s disease patients [48].

We show a connection between FD and co-translational chaperone engagement and dependence. In particular, we show that Ssb, a co-translational chaperone in *S. cerevisiae*, preferentially targets regions of high FD (**Figure 3**). The authors who produced the Ssb data we analyzed here found that in an *in vitro* peptide array – where there is no potential of folding – Ssb recognizes more binding sites than it does *in vivo,* suggesting that some interaction sites are skipped *in vivo* [49]. The authors attribute this discrepancy to additional regulation by cochaperones *in vivo.* However, FD offers an alternative explanation: the skipped sites are simply not exposed long enough. In support of this, we show that the lifetime of the Ssb-Nascent chain complex correlates directly to the FD of the bound segment (**Figure 3G and H**) and that predicted chaperone binding sites that do not match with those found *in vivo* have lower FDs (**Figure 3D and E**). Our proteome-wide analyses suggest that Ssb engagement to high FD regions is not merely opportunistic in the sense that Ssb binds simply because it has the time window to do so. We show that deletion of the Ssb chaperone or chemically disrupting translation disproportionately causes aggregation of proteins with high FDs (**Figure 4A and F**). In fact, proteins that aggregate upon Ssb deletion not only have higher FDs around the Ssb binding site, but they often harbor predicted APRs in exactly those regions (**Figure 4D**). This observation suggests again that Fold-delayed APRs may lead to co-translational aggregation or to off-pathway events that eventually result in protein aggregates, which are mitigated by Ssb. Moreover, proteins of high FDs are also more likely to be co-translationally ubiquitinated, indicating they tend to misfold more readily during translation, and are recognized as such by the cellular machinery (**Figure 4G**).

FD merges two important concepts: fold complexity and ribosome processivity. The latter varies vastly between organisms and conditions. One could argue that a slower translation rate allows for the successful co-translational folding of more complex structures. In support of this, it was shown decades ago that eukaryotic translation systems more efficiently produce modular proteins than their prokaryotic counterparts [50]. However, our FD analyses show that this comes at a cost: slower translation rates mean higher FDs, even for proteins with similar folds (e.g. **Figure 1**). Hence, while slowing down translation opened the door to more complex folds, it may have also necessitated the co-evolution of a more elaborate network of co-translational chaperones because of the associated increase in FD. This may be one of the reasons for the existence of a much more extensive co-translational chaperoning system in eukaryotes than in prokaryotes [51].

Our analyses show that FD can be substantial and have dire consequences. A potential way of evolutionarily reducing FD is codon optimization between long-range interactors to reduce the actual waiting times incurred. Despite substantial efforts, however, we were unable to link FD to codon optimization. We therefore reasoned that perhaps the gain to be made by codon optimization is negligible. Indeed, full optimization of codons theoretically reduces FD by 20% (**Figure 5**). This still leaves substantial FDs, which likely impair co-translational folding. Therefore, we hypothesize that the main strategy for mitigating FD is likely not sequence optimization but chaperone co-evolution, as highlighted by the strong links between Ssb engagement and FD. We suspect that, with the emergence of more data describing *in vivo* binding sites of co-translational chaperones, these will strongly correlate to our FD parameter.

### Limitations of the study

FD, much like CO, is conceptually straightforward. Contacts are simply defined through spatial proximity in the native structure, and it is assumed that folding is delayed until all native interaction partners become physically available. In its current form, the FD algorithm does not consider interactions with the ribosome [52] or intermediate structures, nor does it evaluate the relative contribution of each interaction. Arguably, a residue could adopt its native conformation with only a subset of its interaction partners available. A second limitation of the study is using average codon translation rates to investigate codon optimization. The decoding time of a codon depends on its context within a transcript and has been seen to be condition-dependent [53, 54]. Despite these limitations, FD already offers key biological insights. Moreover, the simplicity of the model makes it computationally inexpensive, allowing us to readily analyze entire proteomes.

## METHODS

### Protein folding vs protein translation rates

Protein folding rates were retrieved from the Protein Folding Database [27] (PFD2.0). This curated dataset contains folding rates derived from experimental data. From the reported folding rates at 25 degrees C (k_f,_), we calculated average folding times (calculated as 1/k_f_). For an estimation of the translation times, proteins from prokaryotic organisms were assigned translation rates of 20 aas/s, whereas proteins form eukaryotes were assigned translation rates of 5 aas/s [4–6]. For an estimation of the total translation time of a protein, we simply multiplied these translation rates by the number of residues in each protein studied.

### SCOPe40 analysis

We analyzed FD profiles of protein domains in the SCOPe40 dataset. This dataset contains single-domain structures that have been manually classified based on their architectures and filtered so that no two domains in the set have more than 40% identical sequences [30, 31]. To establish a general pattern of FD from N– to C-term within domains (**Figure 1I**), residues were assigned relative positions by dividing their position in the domain by the domain length, multiplying by 100 and rounding off to the nearest integer. For each relative position, average FDs and standard deviations were calculated. The average (or mean) FD for a domain was calculated as the sum of the FD of all residues in a domain divided by the domain length.

### Proteome-wide analyses

AlphaFold structures (version 4) and their corresponding predicted aligned error (PAE) matrices for the full proteomes of *Escherichia coli* and *Saccharomyces cerevisiae* (yeast) were retrieved from the AlphaFold Protein Structure Database [32, 33]. Genomic sequences for both species were retrieved from NCBI Genomes FTP server. AlphaFold structures were mapped to genomic sequences using the UniProt ID mapping tool. 3,929 and 4,363 proteins were successfully matched with their corresponding codon sequences for *E. coli* and yeast, respectively. The energies of the structures were minimized using the FoldX “RepairPDB” command, and stability calculations for each amino acid were performed using the “SequenceDetail” command [55]. Protein secondary structures and absolute solvent accessibility values were obtained with DSSP based on the AlphaFold structures [56, 57]. Then, the relative solvent accessibility (RSA) values were calculated by dividing the absolute solvent accessibility values by residue-specific maximal accessibility values, as extracted from Tien *et al* [58]. Disordered regions were defined using the pLDDT score provided in the AlphaFold models, as regions with low confidence scores (pLDDT < 50) have been shown to overlap largely with intrinsically disorder regions [59]. To exclude biases arising from intrinsically disordered proteins, proteins with more than 90% disordered residues were filtered out of the data. Aggregation prone regions were defined with the TANGO algorithm (score > 5) [60] at physiological conditions (pH at 7.5, temperature at 298 K, protein concentration at 1 mM, and ionic strength at 0.15 M).

### FoldDelay

FoldDelay (FD) profiles were determined from protein structures for all SCOPe40 domains and the *E. coli* and yeast proteomes based on AlphaFold models using the formulas described in the Results section. Residues were considered to interact if they contained non-hydrogen atoms within 6 Å. This threshold was chosen since it is commonly used to calculate other topological parameters, such as contact order [3]. For SCOPe40 domains, all residue interactions were considered as the structures were solved with experimental methods. Instead, for AlphaFold predicted models, interactions between two residues whose relative position to each other is low based on the Predicted Aligned Error (PAE) metric were filtered out. Specifically, we excluded interactions with an expected position error > 6 Å.

We assigned a single FD value to each protein to facilitate the proteome-wide FD correlations in **Figure 1** and **Figure 4**. The “mean FD” values correspond to the mean of the FD of individual residues in a structure. The “total FD” values reported are simply the sum of the FD of individual residues in a structure. These metrics provide a global view of the delay incurred by a polypeptide chain throughout its ribosomal production.

### Ssb binding footprints metagene analyses

Ssb binding footprints were obtained from Döring et al. [24] and Stein et al. [25]. Nucleotide positions were transformed to amino acid positions by dividing them by three and rounding down. The lifetime of the Ssb-Nascent chain complex (engagement times) was extrapolated from the width of Ssb binding peaks. Specifically, only Ssb binding peaks with widths falling between 5 and 11 aas were selected for analysis. This range was chosen because higher widths might suggest additional binding and release cycles. A metagene analysis of the Ssb binding footprints was done by aligning the starting site of Ssb binding footprints. The FD profile and the relative enrichment of different properties were calculated, across a range of –120 and 120 aas from the starting site of the Ssb footprints, per position using a rolling average of 3 and after removing empty positions.

As a control measure, random positions were sampled from the same proteins containing the Ssb footprints and the metagene analyses were repeated but aligning on these random positions.

### Comparison between Ssb sites and Limbo regions

Predicted Hsp70 binding regions, here referred to as Limbo regions, were identified with the computational tool Limbo (score > 5) [39]. Döring et al. [24] Ssb binding footprints with a width ranging from 5 to 11 aas were then compared to the Limbo regions. A Limbo region was considered to overlap with an Ssb site if it fell within a range of –55 aas from the starting residue to –35 aas from the ending residue of the Ssb footprints. Based on this criterion, Limbo regions were classified as either “Ssb binding” if there was an overlap or “No Ssb binding” if there was no overlap with any Ssb binding footprint. The difference in FoldDelay between these two groups was obtained by subtracting the median FD values of each group.

### FD of proteins aggregating in Ssb knockout strain

We reanalyzed a dataset produced by Willmund et al. [23]. Through pulldowns of Ribosome Nascent Chain complexes followed by MS, the authors established the *S. cerevisiae* “translatome”. Through Ssb pulldowns, the translatome was then stratified into a group that interacts with Ssb co-translationally (“Ssb not bound” in **Figure 4A**), and a group that does not. The authors further determined which proteins aggregate upon deletion of the Ssb chaperone, indicating they are dependent on Ssb for their solubility. Using this information, we divided the group of Ssb binders into a “soluble” and an “aggregated” fraction as shown in **Figure 4A**.

### FD of proteins sensitive to Arsenite stress

Ibstedt et al. report the identification of aggregated proteins in *S. Cerevisiae* both in physiological conditions (“Physiological” in **Figure 4F**), as well as upon exposure to Arsenite stress (“Arsenic” in **Figure 4F**) [41]. Aggregated fractions were separated through centrifugation and proteins in the aggregated fraction identified through LC-MS. As a background, the authors used a previously established *S. cerevisiae* proteome, which we copied (“MS proteome” in **Figure 4F**).

### FD of proteins that are co-translationally ubiquitinated

Duttler et al. produced a dataset of proteins that are co-translationally ubiquitinated under physiological conditions in *S. cerevisiae* [43]. They do not report a background proteome, so we compared total FD of the co-translationally ubiquitinated proteins with the translatome reported by Willmund et al [23].

### FD optimization through decoding times

To assess whether codon optimization could alleviate FD, we recalculated it for *S. cerevisiae* using codon-specific decoding times as reported by Tuller et al [44], as well as by Sharma et al [46], the latter of which was based on data reported originally by Weismann et al [61]. For each protein, the maxFD was calculated as the furthest interaction in amino acids. To calculate the “actual” FDs reported in **Figure 5**, each codon was assigned its mean decoding time. To calculate “minimal” FD, each codon was assigned the minimal mean decoding time of the codons that encode the same amino acid as the original. Gain was calculated as the difference between the actual FD (s) and the minimal FD (s), representing by how much time FD could in theory be reduced by optimization of codons. The proportional differences were calculated as the gain (s) divided by the actual FD (s).

### Statistics

GraphPad prism or R software were used to perform the different statistical tests. The tests used in each analysis are specified in the corresponding figure. *P*-values are represented as: * *P*-value ≤ 0.05, ** *P*-value ≤ 0.01, *** *P*-value ≤ 0.001 and **** *P*-value ≤ 0.0001.

### Visualizations

Visualizations were performed with GraphPad prism or custom R scripts using the packages ggplot2 [62]. Contact maps were visualized using the circlize R package [63]. ChimeraX was used to visualize protein structures [64].

## ABBREVIATIONS

Aa: Amino acid
CO: Contact order
FD: FoldDelay
APR: Aggregation-prone region

## Funding

The Switch Laboratory was supported by the Flanders Institute for Biotechnology (VIB, grant no. C0401) and the Fund for Scientific Research Flanders (FWO, project grants G053420N and G0A6724N to J.S. and postdoctoral fellowship 12S3722N to B.H.).

## Author contributions

FR and JS conceived and supervised this study. RDR, BH and PFM designed and performed the computational analyses. BH, RDR, FR and JS wrote the manuscript. All authors proofread and corrected the manuscript.

## Competing interests

Subject matter of this publication is part of a patent application (EP 24154112.7) with inventors FR and JS.

## Data and Materials Availability

All data needed to evaluate the conclusions in the paper are present in the paper and/or the Supplementary Materials. The code of all custom R scripts used for data analysis are available from the corresponding authors upon reasonable request.

## Notes

### Competing Interest Statement

Subject matter of this publication is part of a patent application (EP 24154112.7) with inventors Frederic Rousseau and Joost Schymkowitz.

