## Supplementary materials for "Analyzing the implications of protein folding delay caused by translation"

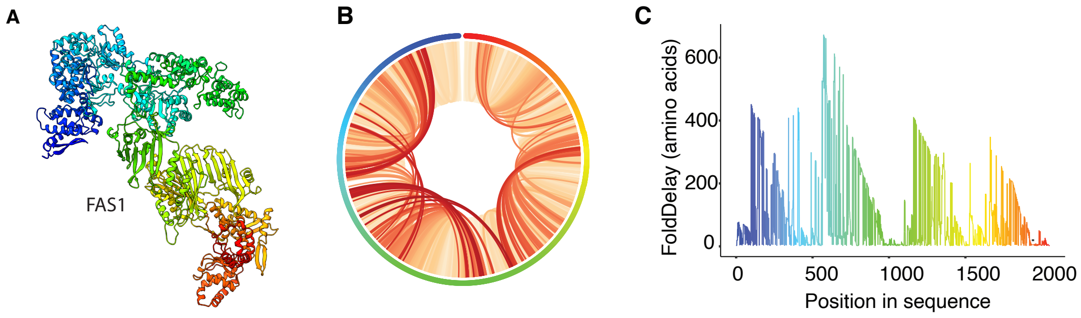


**Supplementary Figure 1 – Multidomain proteins display a sawtooth FD profile. (A)** AlphaFold-predicted structure of the yeast multidomain protein Fatty acid synthase subunit beta (FAS1; UniProt ID P07149). **(B)** Contact map derived from the structure in (A). **(C)** FoldDelay profile of FAS1.

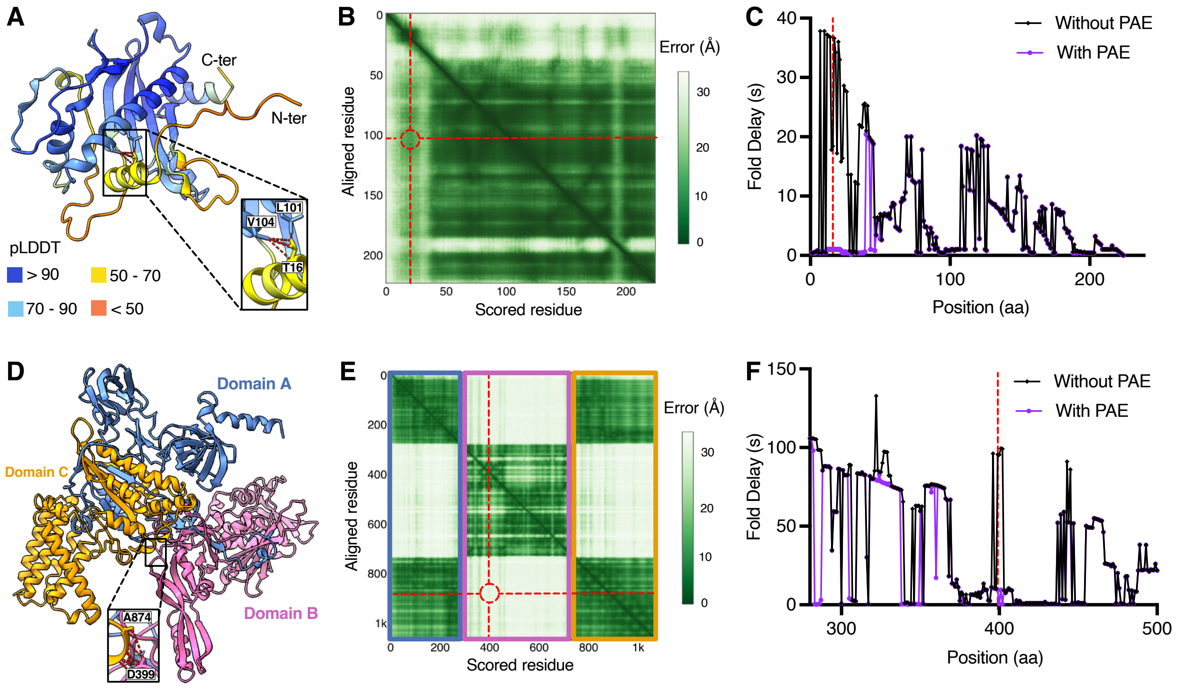


**Supplementary Figure 2 – Predicted aligned error (PAE) to assess the reliability of residue interactions. (A)** The N-terminal region of the yeast protein Holocytochrome-c1 synthase (UniProt ID: Q00873) is structurally disordered (low pLDDT scores). AlphaFold randomly places this region in space, which leads to some interactions that are not real, such as the interactions between residue T16 and L101/V104. **(B)** PAE plot for Holocytochrome-c1 synthase. The colour at (x, y) corresponds to the expected distance error in the residue x's position when the predicted and the true structures are aligned on residue y. The expected distance error between residue T16 and residues L101/V104 is shown in red. **(C)** FD profiles for Holocytochrome-c1 synthase before and after filtering interactions between residues with an expected distance error > 6Å. Residue T16 is highlighted in red. **(D)** AlphaFold model for the yeast protein V-ATPase subunit A (UniProt ID: P17255). Domains are shown in different colors. The interaction between residue D399 in domain B and residue A874 in domain C is highlighted. **(E)** PAE plot for V-ATPase subunit A. The relative orientation of domain B with domains A and C has a very low confidence. Therefore, the interaction between D399 and residue A874 (shown in red) will probably not occur in the protein. **(F)** FD profiles for V-ATPase subunit A before and after filtering interactions using the same threshold as (C). Residue D399 is highlighted in red.

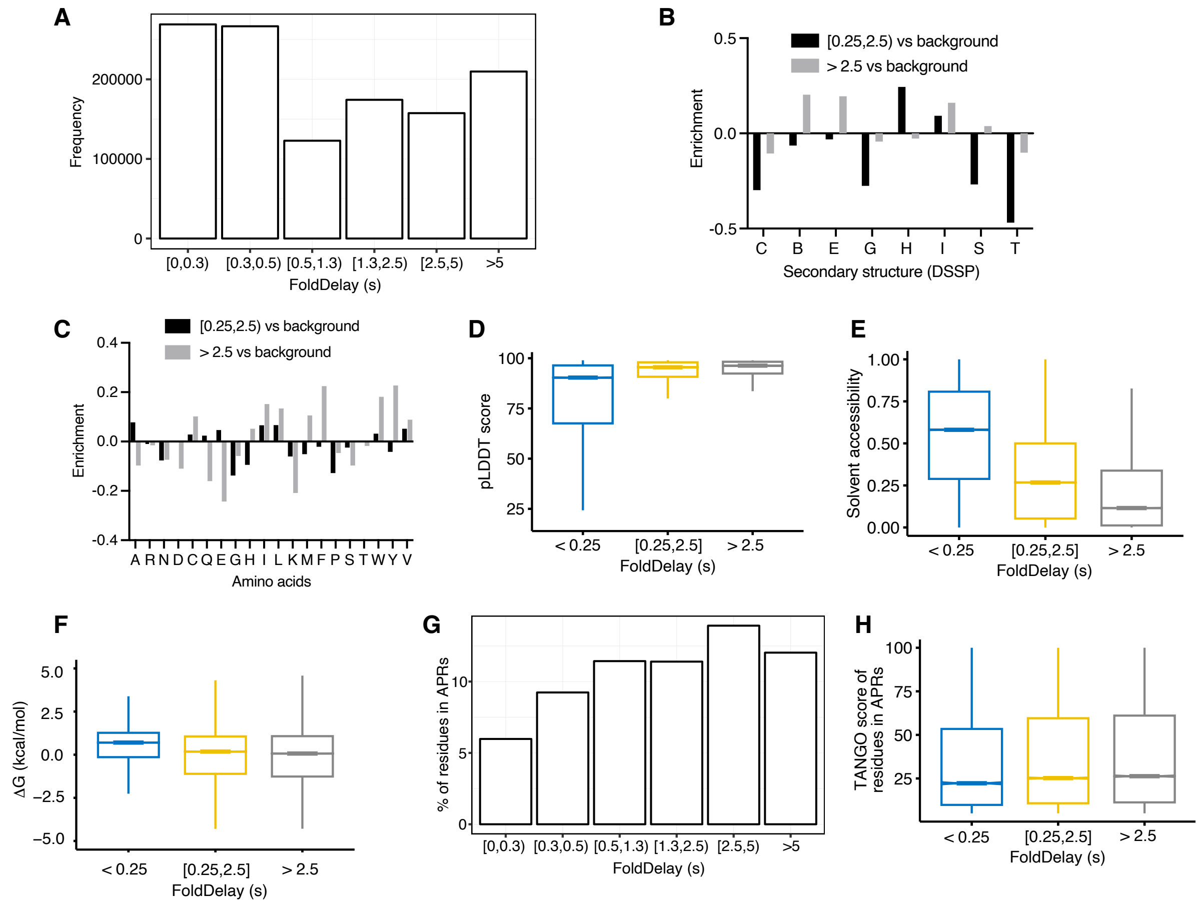


**Supplementary Figure 3 – Fold Delay captures the vectorial nature of protein translation.** **(A)** Number of residues for different FD bins in *E. coli* proteins. **(B)** Enrichment of residues with FDs between 0.25 and 2.5 seconds (or bigger than 2.5 seconds) versus background for the different DSSP secondary structure categories in *E. coli* proteins. C = coil, B = β-bridge, E = extended strand in β-sheet conformation, G= 3-turn helix, H = 4-turn helix, I = 5-turn helix, S= bend and T = hydrogen bounded turn. **(C)** Enrichment of residues with FD between 0.25 and 2.5 seconds (or bigger than 2.5 seconds) versus background for all amino acid types in *E. coli* proteins. **(D-F)** pLDDT scores (E), solvent accessibilities (F) and stabilities for residues in yeast proteins for different categories of FD. **(G)** Percentage of residues in APRs (TANGO score > 5) for different FD bins in *E. coli* proteins. Residues in transmembrane domains and signal peptides were filtered out to avoid biases. **(H)** Aggregation strength (TANGO score) for residues in APRs of *E. coli* proteins for different categories of FD.

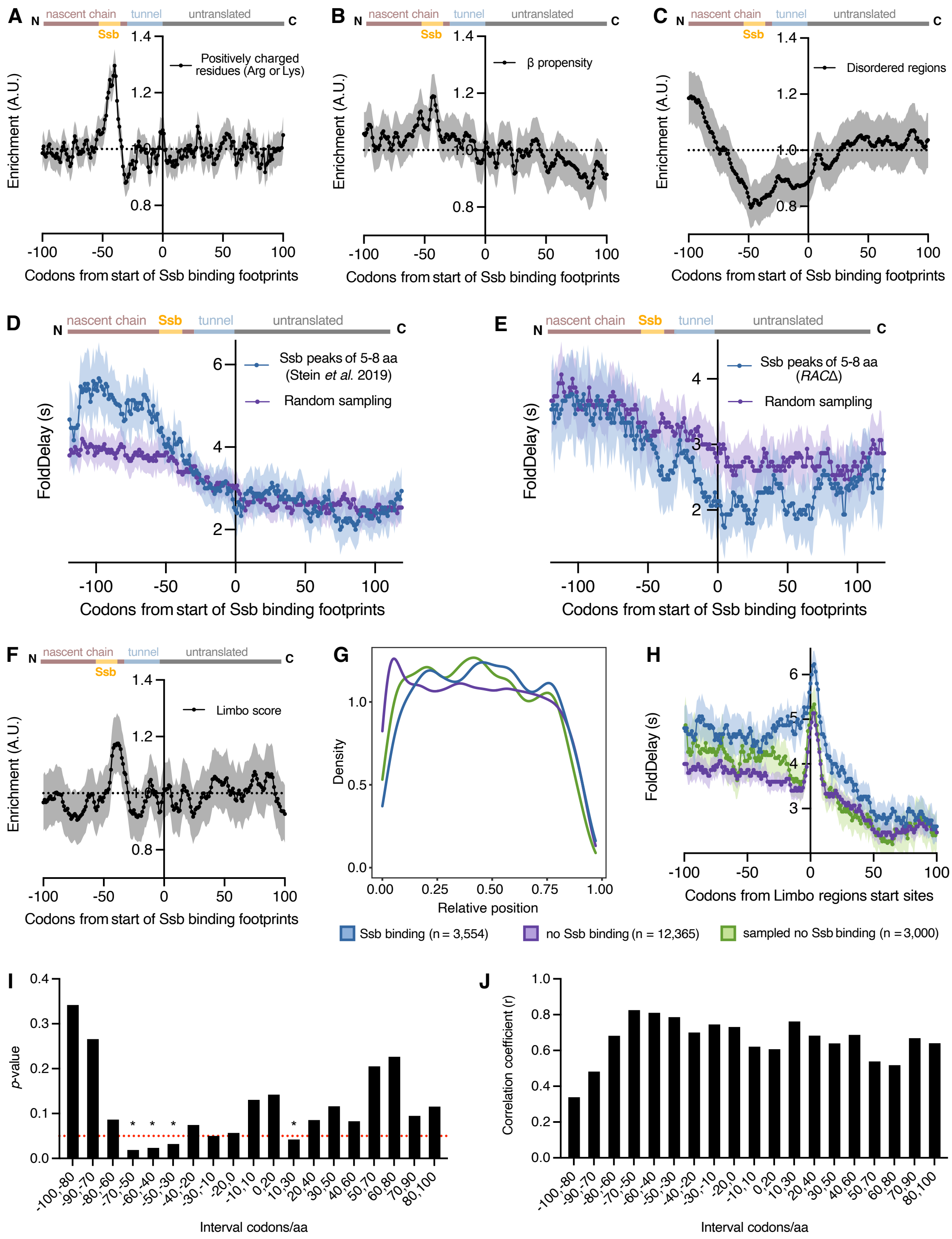


**Supplementary Figure 4 – Ssb binds to regions with high Fold Delays. (A-C)** Relative enrichment of the number of positively charged residues (A), β-propensity (B) and structural disorder (C) in the nascent chain at the start of Ssb binding for sites with a peak width between 6-8 aa (n = 3,371). The line represents the median value at each position, while the shaded region is the 95% bootstrapped confidence interval (CI). **(D and E)** Fold delay in the nascent chain at the start of Ssb binding for sites with a peak width between 5-8 aa (n = 1,798) from Stein et al. (*1*) (D) and for sites in *RAC*Δ cells with a peak width between 5-8 aa (n = 1,571) (E), as compared to 4,000 randomly sampled positions from the same sets of proteins. The line represents the median value at each position, while the shaded region is the 95% bootstrapped CI. **(F)** Relative enrichment of Limbo scores in the nascent chain at the start of Ssb binding for sites with a peak width between 6-8 aa (n = 3,371). The line represents the median value at each position, while the shaded region is the 95% bootstrapped CI. **(G)** Density distributions showing the relative positions of Limbo regions that are Ssb binding sites (blue), not Ssb binding sites (purple) and not Ssb binding sites that have been sampled to follow the same distribution as Ssb binding sites. **(H)** Fold delay of the same set of regions as in (G). The line represents the median value at each position, while the shaded region is the 95% bootstrapped CI. (**I and J)** *P*-values (I) and correlation coefficients (J) of the average median FDs per width peak at different interval positions of aligned Ssb binding footprints with a peak width between 5-11 aa. *P*-values lower than 0.05 (indicated by a red line) are shown with an asterisk. The average FD value increases with Ssb peak width only close to the Ssb binding region. Experimental Ssb binding sites used in all panels (except D) are derived from (*2*).

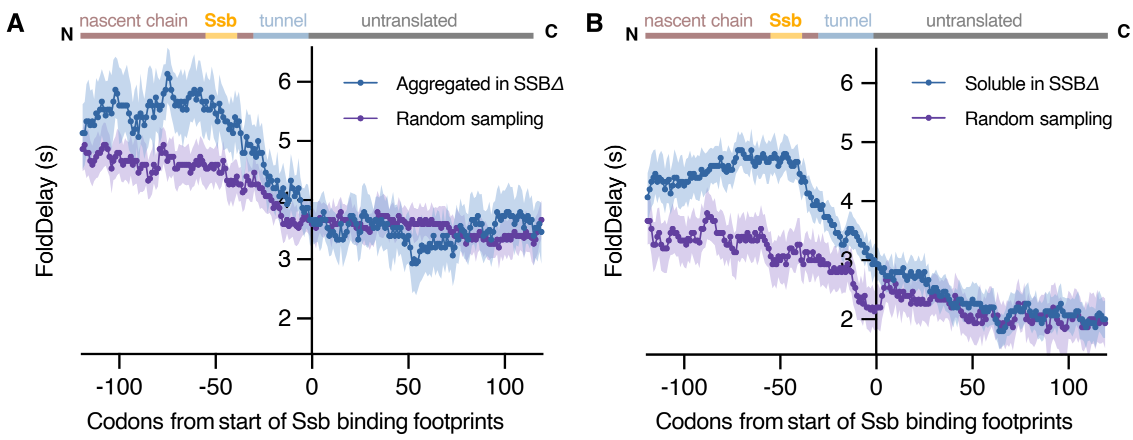


**Supplementary Figure 5 – Proteins with high Fold Delays are associated with co-translational misfolding and aggregation. (A)** Fold delay in the nascent chain at the start of Ssb binding for sites with a peak width between 5-11 aa (n = 1,917) in proteins that aggregate upon deletion of Ssb as compared to 4,000 randomly sampled positions from the same set of proteins. **(B)** Fold delay in the nascent chain at the start of Ssb binding for sites with a peak width between 5-11 aa (n = 5,415) in proteins that do not aggregate upon deletion of Ssb as compared to 4,000 randomly sampled positions from the same set of proteins. The line represents the median value at each position, while the shaded region is the 95% bootstrapped CI (A and B). Experimental Ssb binding sites used in both panels are derived from (*2*).
